# Tubulin isotypes polymerise into sectioned microtubules that locally regulate protein binding

**DOI:** 10.1101/2025.03.12.642813

**Authors:** Manjari Prakash, Yean Ming Chew, Denisa Ondruskova, Jiri Miksatko, Alica Dodokova, Arya Krishnan, Carsten Janke, Robert A. Cross, Zdenek Lansky, Marcus Braun

## Abstract

Microtubules assemble from tubulin heterodimers composed of conserved α- and β- tubulins. In many species, including humans, tubulins are expressed from multiple genes. While the resulting tubulin isotypes–show only subtle sequence differences, microtubules made of distinct isotypes can differ in their dynamic behaviour, as well as in their structure, for example in the number of their protofilaments. In cells, tubulin isotypes co-polymerize into mixed-isotype microtubules. How the mixing of tubulin isotypes affects microtubule functionality is unknown. Here we show that co-polymerization of recombinant tubulin dimers containing two different human β-tubulin isotypes, α1β3 and α1β4, generates sectioned microtubules in which tubulin content and protofilament number differ from one section to the next. We demonstrate that two microtubule-associated proteins (MAPs), TPPP1 and optineurin (OPTN), bind differentially to these sections. Microtubules grown from natively mixed-isotype sources, either HeLa cells or mammalian brain tissue, also consist of sections that are differentially recognized by TPPP1 and OPTN, suggestive of isotype sorting. Our results demonstrate that co-polymerization of multiple tubulin isotypes can drive microtubules to assemble into distinct sections that locally regulate the interactions of MAPs with the microtubule lattice.

## Introduction

Microtubules play important roles in cells, maintaining structural stability, serving as tracks for motor proteins or facilitating cell division. Microtubules are polymers composed of heterodimers of α- and β-tubulin (Mitchison & Kirschner, 1984; Nogales, 2000). They polymerize in a linear array to form protofilaments, which associate laterally forming a hollow tube (Kalutskii et al., 2024). In vitro polymerized microtubules can consist of 11 to 16 protofilaments (Chaaban & Brouhard, 2017) and the number of protofilaments can change within a single microtubule (Chrétien et al., 1992; Guyomar et al., 2022). By contrast, the number of protofilaments in cellular microtubules is mostly 13 (Tilney et al., 1973), however variations have been reported in several model organisms due to a more pronounced tubulin isotype heterogeneity. Evidence from nematode *Caenorhabditis elegans* show the touch receptor neurons have a cell-specific protofilament number of 15 due to the tubulin isotypes mec-7 and mec-12. This specific protofilament number of 15 in these neurons is essential for the touch-sensing function of these neurons (Savage et al., 1989). Mammalian cells also can have distinct protofilament microtubules, for example, microtubules in sensory cell of the mammalian inner ear also have a protofilament number of 15 (Kikuchi, Takasaka, Tonosaki, Katori, et al., 1991; Kikuchi, Takasaka, Tonosaki, Watanabe, et al., 1991; Renauld et al., 2021).

Humans have 9 α-tubulin and 10 β-tubulin isotypes (Nsamba & Gupta, 2022). These isotypes have varied expression and distribution across different cell types and tissues. Additionally, the C– terminal tails of α- and β-tubulins are subjected to a range of post-translational modifications. Both, tubulin post-translational modification and tubulin isotypes are expected to regulate the interaction of microtubule-associated proteins and molecular motors with microtubules (Janke & Magiera, 2020), however very little is so far known about the role of isotypes. Microtubules formed *in vitro* using single recombinant tubulin isotypes have been shown to adapt distinctive in structures and dynamic behaviours (Chew & Cross, 2023; Ti et al., 2018; Vemu et al., 2017). Human α1β3-tubulin *in vitro* predominantly assembles into 13-protofilament microtubules, which are more dynamic than the predominantly 14 protofilament microtubules assembled from recombinant α1β2-tubulin (Ti et al., 2018). The number of protofilaments in insect sperm microtubules varies between species. Cross-expression of specific tubulin isotypes in different insect species shows that the protofilament number of the assemble microtubules is a property intrinsic to these specific tubulin isotopes (Raff et al., 1997).

Several *in vitro* studies have shown that the dynamic behaviour of microtubules is a function of isotype composition. Adding recombinant α1β3-tubulin to mixtures of tubulin derived from sources that natively do not contain any α1β3 increases the dynamics of the assembled microtubules (Vemu et al., 2017). Tubulin isotype composition can also differentially regulate microtubule-associated proteins, such as the kinesin-related protein MCAK. Microtubules formed from α1β3 are more susceptible to MCAK-driven depolymerization than α1β2 microtubules. Microtubules formed from a mixture of α1β3 and α1β2 tubulins have intermediate stability during MCAK-dependent depolymerization, suggesting a dose-dependency of the isotype-related effects (Ti et al., 2018). Similarly, intermixing of α1β3 and α1β4 in various ratios can differentially tune the interaction of kinesin-1 molecular motor with the microtubules (Chew & Cross, 2023). Given these functional differences of microtubules made from different isotypes, and exciting question is how tubulin isotypes co-assemble into microtubules – do they form a mixed lattice, or can they generate discrete regions that are functionally different, for instance because of differential regulation of microtubule-associated proteins.

Here we show that the microtubule-associated protein OPTN (Liu et al., 2025) shows a preference for microtubules assembled from α1β4-tubulin over α1β3-microtubules, as well as a preference for 14- over 13 or 12-protofilament microtubules. TPPP1, another microtubule-associated protein (Schofield & Bernard, 2013), did also preferentially bind α1β4-microtubules, however this binding is not affected by the protofilament number. When we copolymerized microtubules from α1β3- and α1β4-tubulin, we observed that OPTN and TPPP1 segregate into distinct sections of the microtubules, which we show correspond to microtubule sections of distinct protofilament numbers and with distinct isotype composition. Furthermore, we show that the microtubules grown *in vitro* from native mixtures of tubulin isotypes, purified either from HeLa cells or from mammalian brain tissue, equally form sections that are selectively recognized by TPPP1 or OPTN. These findings demonstrate that tubulin isotype mixtures do not assemble uniformly into homogenous microtubules but form sectioned microtubules in which the sections interact differently with specific MAPs.

## Results

### TPPP1 and OPTN differentially interact with microtubules polymerized from isotype α1β4 and α1β3

The tubulin isotypes α1β4 and α1β3 differ in their expression in different cell types, with α1β4 showing a wider distribution across many cell types whilst α1β3 is more specific to neurons (Banerjee et al., 1988). To check for a difference in MAP binding to these to tubulin isotypes, we polymerized single-isotype microtubules from recombinant, purified α1β4 and α1β3 tubulin (Chew & Cross, 2023) with GTP. Polymerised microtubules were stabilised with taxol. We then immobilized a first type of microtubules in a flow channel, imaged them using IRM (interference reflection microscopy, Methods), and subsequently added the second microtubule type followed by IRM imaging (Figure 1A). This allowed us to distinguish between two sets of unlabelled microtubules. Finally, we added fluorescently labelled MAPs, either in a lysate or as purified protein, to the microtubules (Methods). Out of a pool of tested eight MAPs, we identified TPPP1 and OPTN as differentially binding to α1β4 versus α1β3 single-isotype microtubules. We found that both TPPP1 and OPTN preferentially bind to microtubules made of α1β4 as compared to α1β3 tubulin (Figure 1B,C). In the presence of taxol, α1β4 microtubules adopt an expanded lattice, while α1β3 microtubules remain in their compacted form (Chew & Cross, 2023). To test if lattice expansion affects the binding of TPPP1 and OPTN we polymerized the α1β4 or α1β3 microtubules in presence of GMPCPP, the non-hydrolysable analogue of GTP, which for microtubules polymerized from mixed-isotype brain tubulin results in extended microtubules (Hyman et al., 1995). We find that for GMPCPP lattices also, both TPPP1 and OPTN preferentially bind to α1β4 microtubules as compared to α1β3 (Fig. S1A,B,C). To test the binding of TPPP1 and OPTN to natively compacted GDP lattices, unperturbed by microtubule-targeting drugs or nonhydrolyzable nucleotides, we attenuated microtubule depolymerization by addition of glycerol to the buffer (Keates, 1980). Also under these conditions, both TPPP1 and OPTN preferentially interact with α1β4 compared to α1β3 microtubules (Fig. 1C,D). We conclude that both TPPP1 and OPTN differentiate between single-isotype microtubules polymerized from α1β4 or α1β3 tubulins, independent of tubulin nucleotide states, microtubule lattice expansion, and the interaction with the stabilizing drug taxol. Hence TPPP1 and OPTN can act as probes for tubulin isotype composition.

**Figure 1:**
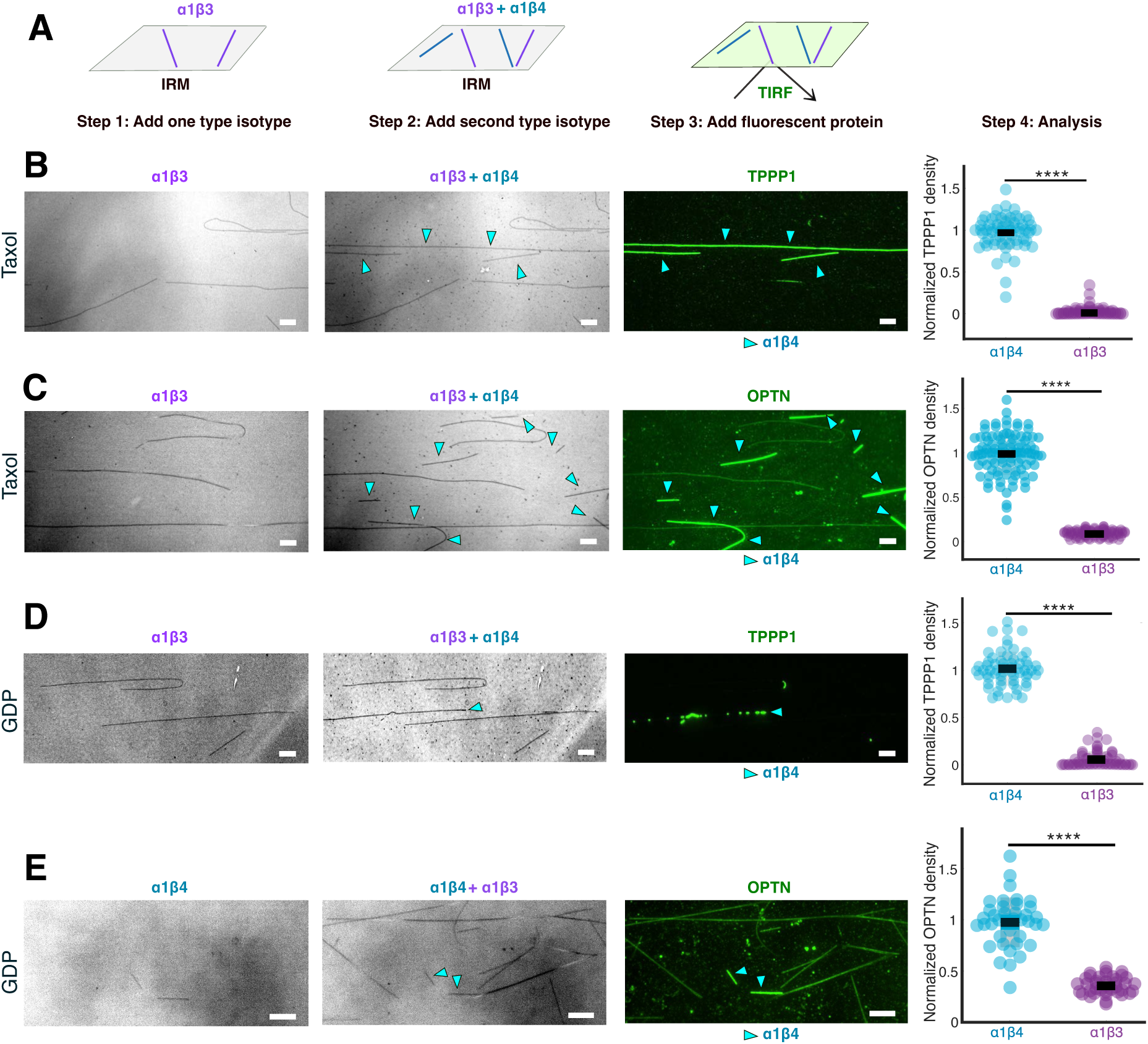
TPPP1 and OPTN differentially interact with microtubules polymerized from isotype α1β4 and α1β3. A) Experimental setup. Sequential flushes of tubulin isotypes imaged by IRM, allowing discrimination of isotypes, after which fluorescent OPTN or TPPP1 is added, imaged using TIRF microscopy and analysed. B,C) Taxol-stabilized α1β3 and α1β4 single-isotype-microtubules are sequentially flushed in and later B) TPPP1 or C) OPTN is imaged, and fluorescence density quantified (right). D,E) GDP lattice α1β3 and α1β4 single isotype microtubules are sequentially flushed in and imaged by IRM. Later D) TPPP1 or E) OPTN is imaged, and fluorescence density is quantified from the whole microtubule (right). Cyan arrows point to the α1β4 single isotype. Scale bars 4 µm. N= 3 for each experiment in figure, **** is p<0.0001, Student t test for each plot in figure.

### Mixtures of α1β3 and α1β4 tubulin assemble into sectioned microtubules of locally distinct composition

Given the different dynamics of α1β3 and α1β4 microtubules (Chew & Cross, 2023), we wondered how these tubulin isotypes would intermix when forming microtubule lattices by co-polymerization, and how these intermixed lattices might interact with TPPP1 and OPTN. We polymerized microtubules from mixtures of α1β3 and α1β4 tubulin in presence of GTP in the ratio of 1:2 of α1β3 to α1β4, followed by taxol addition to stabilize the formed microtubules. α1β3 and α1β4 microtubules were then immobilized on a coverslip. We next added OPTN, which was subsequently washed out and replaced by TPPP1. We found that the distributions of both OPTN and TPPP1 on these mixed-isotype microtubules were non-uniform, with the two MAPs having higher binding occupancy on certain sections of the microtubule lattice (Figure 2A), with both OPTN and TPPP1 preferring the same sections. This indicated that the mixture of purified α1β3 and α1β4 tubulin might have self-assembled into sectioned microtubules of locally distinct composition.

**Figure 2:**
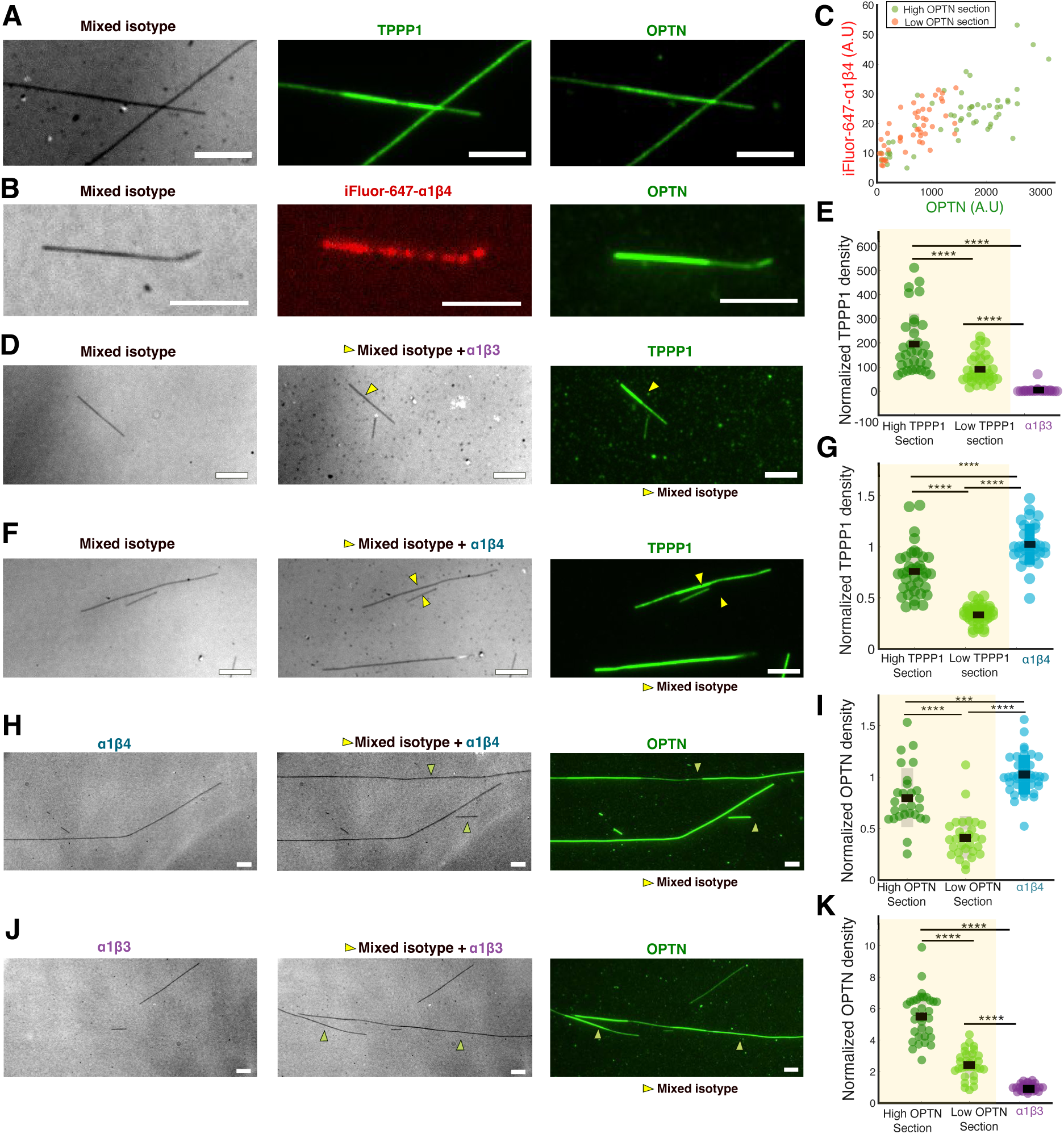
Mixtures of tubulin isotypes α1β3 and α1β4 assemble into sectioned microtubules of locally distinct composition. A) IRM image of mixed isotype microtubules, TPPP1 on the microtubule, after removing TPPP1, OPTN imaged on the same microtubule. B) IRM image of mixed isotype microtubules, iFluor-647-α1β4 tubulin channel, OPTN on the microtubule. C) Correlation plot of quantification of iFluor-647-α1β4 signal and OPTN from the same regions on identified high-density section and low-density sections, Pearson correlation coefficient is 0.716, r² is 0.512, p<0.001. D) Microtubule interaction of TPPP1 on unlabelled mixed isotype microtubules (α1β3: α1β4 in 1:2 ratio) in comparison with single isotype α1β3, sequential IRM images of the microtubules flushed in, TPPP1 on microtubules, mixed isotypes marked with yellow arrows. E) Analysis of TPPP1 signal on the high-density section, low-density section of mixed isotype microtubules compared with α1β3. F) Microtubule interaction of TPPP1 on unlabelled mixed isotype microtubules (α1β3: α1β4 in 1:2 ratio) in comparison with single isotype α1β4, sequential IRM images of the microtubules flushed in, TPPP1 on microtubules, mixed isotypes marked with yellow arrows. G) Analysis of TPPP1 signal on the high-density section, low density section of mixed isotype microtubules compared with α1β4. H) Microtubule interaction of OPTN on single isotype α1β4 compared with unlabelled mixed isotype microtubules (α1β3: α1β4 in 1:2 ratio), sequential IRM images, OPTN on microtubules. I) Analysis of OPTN signal on the high-density section, low-density section and α1β4. J) Microtubule interaction of OPTN on single isotype α1β3 compared with unlabelled mixed isotype microtubules (α1β3: α1β4 in 1:2 ratio), sequential IRM images, OPTN on microtubules. K) Analysis of OPTN signal on the high-density section, low density section and α1β3. Shaded regions of the plots in yellow indicate mixed isotype microtubules. Arrows on the images indicates mixed isotype microtubules, Scale bars 4 µm. N= 3 for each experiment in figure, **** is p<0.0001, Student t test for each plot in each plot in figure.

Given the previously observed higher binding affinity of OPTN and TPPP1 to α1β4 tubulin (Fig. 1B-E), we hypothesized that the MAP-recognized sections might contain different amounts of α1β4. To directly visualize tubulin isotype distributions in our polymerized microtubules we labelled purified α1β4 tubulin with iFluor-647 (Methods), grew microtubules from mixtures of α1β3 and iFluor-647-α1β4 tubulin, again in the ratio of 1:2 of α1β3 to α1β4, immobilized them on coverslips and then imaged the distribution of the fluorophores and found them to be unevenly distributed (Fig. S1E). Next, we added OPTN to mark the different sections by differential OPTN binding. We binarised the OPTN fluorescent signal on microtubules into high-density and low-density sections, to then compare the iFluor-647-α1β4 tubulin fluorescent signal within these sections. We found that strong iFluor-647-α1β4 correlates with strong OPTN fluorescent signal (Fig. 2B,C; S1E,F). This result suggests that microtubules formed by the α1β3 and α1β4 tubulin mixture self-assembled to distinct sections with different α1β3 and α1β4 tubulin content.

We next compared OPTN and TPPP1 binding on these mixed-isotype microtubules to microtubules polymerized solely from α1β3 and α1β4. We found that the sections recognized by OPTN on mixed-isotype microtubules exhibited lower OPTN fluorescence signal than the the signal measured on single-isotype α1β4 microtubules, while regions of low OPTN binding exhibited higher OPTN fluorescence density than on single-isotype α1β3 microtubules (Fig. 2H-K). Quantifying the OPTN fluorescence in these regions and comparing it to pure a α1β4 or α1β3 microtubules we estimated the tubulin content to be 74% α1β4 in the high OPTN binding regions and 26% α1β4 in the low OPTN binding regions. Using TPPP1 as an indicator of tubulin isotype content, we obtained strikingly similar results: 72% α1β4 in the high-TPPPl binding, and 28% α1β4 in low-TPPPl binding microtubule sections. Combined, these experiments show that microtubules can polymerize into sections that are enriched in specific tubulin isotypes, which in turn differentially regulates the interactions of MAPs with these sections.

### OPTN-bound microtubule sections have higher protofilament numbers

It has been previously shown that the human β-tubulin isotypes can regulate microtubule protofilament numbers: α1β3 tubulin polymerises predominantly into microtubules of 13 protofilaments, while α1β2 tubulin rather forms 14-protofilament microtubules (Ti et al., 2018). So far, however, the protofilament number of microtubules formed from α1β4 tubulin is not known.

To determine whether the protofilament numbers of microtubules assembled from α1β4 tubulin, or a 1:2 mix of α1β3 and α1β4 tubulin isotype have an effect on OPTN or TPPP1 interaction with microtubules, we first tested whether TPPP1 or OPTN are sensitive to protofilament numbers at all. Using tubulin purified from mammalian brain, we polymerized microtubules in two conditions previously described to respectively result in predominantly 12- or 14-protofilament microtubules, (Ray et al., 1993). 12 protofilament microtubules are polymerized by addition of 2 uM Taxol during polymerization, and 14 protofilament microtubules were polymerized by using GMPCPP. After confirming protofilament numbers by electron microscopy (Fig. S2B,D). Interference Reflection Microscopy (IRM) provided a ∼0.2-fold reduction in signal obtained for 12- versus 14- protofilament microtubules (Fig. S2A, C), reflecting the smaller diameter of microtubules with a smaller protofilaments number. When immobilizing the two sets of microtubules in a single imaging chambers and adding fluorescent TPPP1 or OPTN we found that TPPP1 did not distinguish between the two sets of microtubules, while OPTN displayed a ∼2 fold preference for 14-protofilament microtubules (Fig. 3C,D). This demonstrates that OPTN is sensitive to protofilament numbers, while TPPP1 is not. Next, we applied the same method to the α1β3 and α1β4 single-isotype microtubules and observed a ∼0.1 fold reduction (p<0.0001) in IRM signal for α1β3 single-isotype microtubules (Fig. 3E,F), suggesting a difference of about one protofilament. OPTN preferential binding to α1β4-, compared with α1β3-, single-isotype microtubules (Fig. 3F) hence is caused by the combined effect of OPTN higher affinity for (a) microtubules assembled from α1β4-tubulin (as compared with α1β3-tubulin, Fig. 3E,F) as well as (b) a preference for higher protofilament numbers of α1β4-tubulin microtubules as determined by IRM (Fig. 3F).

**Figure 3:**
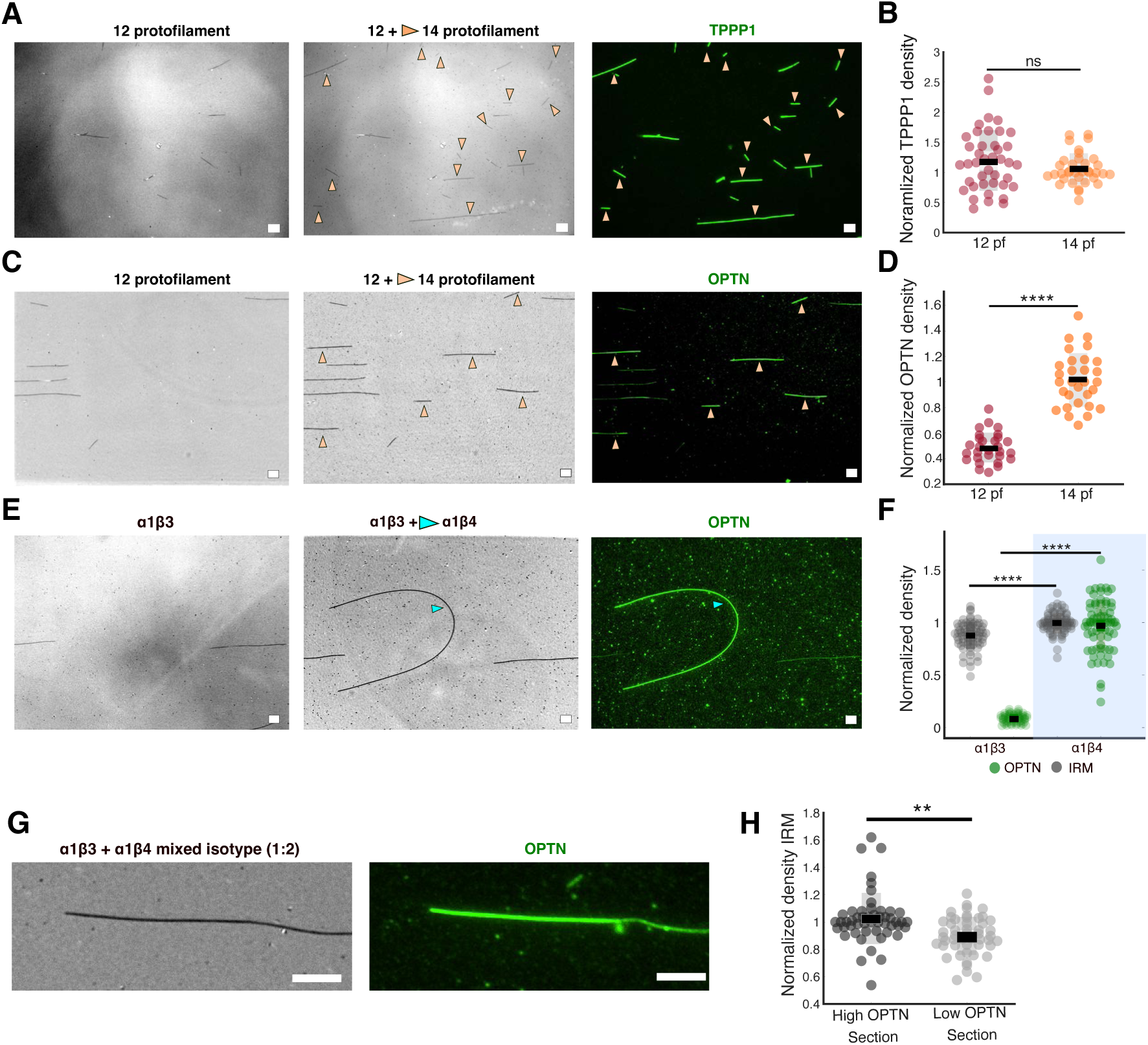
OPTN identifies microtubule sections of higher protofilament numbers. A) IRM images of serial flushes of 12 and 14 protofilament microtubules, TPPP1 on microtubules, orange arrows indicate 14 protofilament microtubules. B) Analysis of TPPP1 signal on 12 and 14 protofilaments, p=0.1392. C) IRM images of serial flushes of 12/14 protofilament microtubules, OPTN imaged on microtubules, orange arrows indicate 14 protofilament microtubules. D) Analysis of OPTN signal on 12 and 14 protofilament number microtubules, p<0.0001. E) IRM images of serial flushes of taxol-stabilized α1β3 followed by α1β4, OPTN on the microtubules, cyan arrow points to α1β4 isotype. F) Analysis of IRM signal of α1β4- and α1β3-microtubules done in parallel, with quantification of OPTN signal on the same microtubules, p<0.0001. G) IRM image of mixed-isotype microtubules, OPTN on mixed isotype microtubules showing high-OPTN and low-OPTN sections. H) Analysis of IRM density on OPTN recognized section and low-OPTN section on mixed-isotype microtubules, ** means p=0.0012, Scale bars 4 μm N= 3 for each experiment in figure. Student t test for each plot in figure. 12 pf: 12 protofilaments, 14 pf: 14 protofilaments.

Having established the IRM method to evaluate the number of protofilaments, we next employed the method to estimate the protofilament numbers in the sections we observed on mixed-isotype microtubules. After having taken IRM images, we added OPTN to identify sections of high-affinity binding of this protein. We found that the IRM signal intensities was about 0.1-fold higher in sections of higher OPTN fluorescence (Fig. 3G-H), suggesting that sections of high α1β4 content have more protofilaments, while sections of low α1β4 content have fewer protofilaments. Combined, these results show that tubulin isotypes, upon co-polymerization *in vitro*, form discrete sections that are enriched in specific tubulin isotypes and have distinct protofilament numbers, which regulates the interaction of MAPs with those sections.

### TPPP1 and OPTN highlight distinct sections on microtubules assembled from natively mixed-isotype tubulin purified from HeLa cells or mammalian brain

In vivo, microtubules are composed of tubulin isotype mixtures that vary across cell types. Additionally, tubulin is subjected to various post-translational modifications. Mammalian tubulin purified from brain tissue, which commonly used for *in vitro* reconstitution assays, is enriched in several known tubulin modifications, while tubulin from HeLa cells has very low to indetectable levels of those modifications (Barisic et al., 2015). To assess the role of tubulin isotypes in the absence of post-translational modifications, we assembled microtubules from HeLa tubulin and added either fluorescent TPPP1 or OPTN. We found both proteins bind the microtubules in regions of two discrete densities (Fig. 4A). Exchanging the proteins in the same imaging chamber revealed that their regions of preferential binding are identical (Fig. 4B, C-E). Similarly, removing TPPP1 or OPTN from the imaging chamber, which resulted in unbinding of the proteins from microtubules, followed by re-introduction of the same proteins into the chambers led to their re-binding to the same microtubule sections (Fig. 4C-E, Movie S1). This experiment indicates that the sections of high binding on the microtubule, likely defined by a specific isotype composition, determine the interaction affinities. When we tested other MAPs, such as CKAP5 (Sabo et al., 2024), a human homolog of XMAP215, we found that, irrespective of the sections recognized by OPTN on microtubules, CKAP5 uniformly bound to the entire microtubules (Fig. 4F). This shows that the sensitivity of TPPP1 and OPTN for these regions is not a general feature of other MAPs. We further found that OPTN also binds to specific sections on the native (glycerol-stabilized) HeLa GDP microtubules, excluding that taxol stabilization is the cause for this behaviour (Fig. S1D). FRAP experiments on HeLa microtubules quantifying the recovery of OPTN in regions of high-density and low-density binding showed a modest difference in recovery time(s) (Fig. 4I, S3A,B). OPTN interacted longer in sections of high binding on HeLa microtubules, which showed higher protofilament numbers as revealed by our IRM method (Fig. 4G, H).

**Figure 4:**
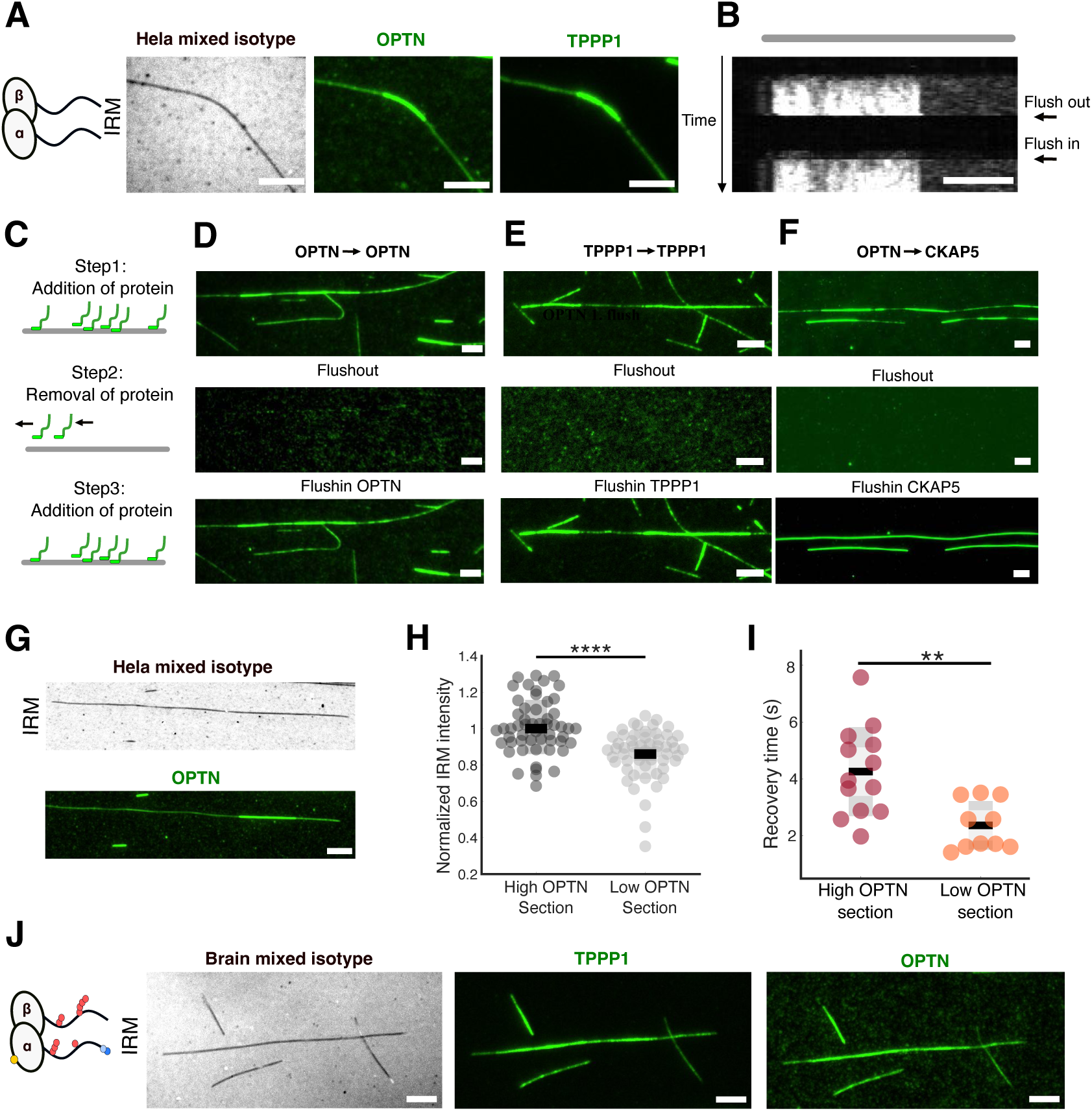
TPPP1 and OPTN distinguish distinct sections on microtubules assembled from native tubulin mixtures purified from HeLa cells or mammalian brain. A) Schematic representation of HeLa tubulin without post-translational modifications, IRM image of a HeLa microtubule, OPTN on microtubules, TPPP1 on the same microtubule after flushing-out of OPTN. B) Kymograph depicting section of TPPP1. flush-out of TPPP1 (44 seconds post start of imaging), flush-in TPPP1 back (77 sec post start of imaging), vertical scale 1.7-min, grey line on the top of the kymograph indicates the length of the microtubule. C) (Top to bottom) Schematic representation of protein - addition, removal and re-addition to the same channel. D.E.F) (Top to bottom) addition of proteins D) OPTN E) TPPP1 F) OPTN, removal of protein, and later flush in of the proteins D) OPTN, E) TPPP1. F) CKAP5. G) (Top to bottom) IRM image of HeLa microtubules after background subtraction. OPTN on the microtubule identifying the high intensity sections and low intensity sections. H) Analysis of IRM density on the high OPTN section and the low OPTN section of the HeLa microtubule. I) Recovery time calculated for the high/low OPTN section based on intensity recovery curves on the HeLa microtubules, p=0.0025, J) Schematic representation of highly post-translational modified brain tubulin, IRM image of a microtubule assembled from brain tubulin, TPPP1 on the microtubule identifying the high-density and low-density sections, OPTN on the microtubule recognising the same high-density and low-density sections. Scale bars 4 µm. N= 3 for each experiment in figure, **** is p<0.0001, ** is p<0.01, Student t test for each plot in each figure.

We finally tested the impact of post-translational modification on the binding behaviour of TPPP1 and OPTN. Repeating the assays with microtubules polymerized from brain tubulin we found both proteins binding non-uniformly, in discrete densities to microtubules (Fig. 4J), albeit not as distinctly different as on HeLa microtubules. We attribute the reduction of the effect to tubulin post-translational modifications masking the preferential binding to tubulin isotypes. Combined, these results suggest that tubulin from native sources can form discrete sections made of specific isotypes, and having distinct protofilament numbers, which can guide the binding of certain of MAPs to those sections. Our findings implies that microtubules can be patterned into distinct sections, which in cells might acquire specialized functions through selective interactions with MAPs.

## Discussion

The tubulin code postulates that biochemical diversity of the building blocks of the microtubules, the tubulin isotopes could determine the behaviour and functions of microtubules. However, whether tubulin isotypes simply co-polymerise into homogeneously mixed microtubules, or whether they can generate patterned microtubules has been so far not understood. The functional consequences of these two different models of co-polymerisation are quite different: while in the first case microtubules would be highly similar in a given cellular environment, in the latter case we could expect to form microtubules of different isotype composition side by side in the same cell, or even patterned microtubules consisting of distinct stretches made of specific tubulin isotypes, and thereby acquiring specific properties and functions. In our current work we have tested the model experimentally and found the differential regulation of MAPs TPPP1 and OPTN via the assembly of mixed tubulin isotypes into sectioned microtubule lattices. TPPP1 shows increased affinity for the isotype α1β4 over α1β3 irrespective of the protofilament numbers, while OPTN in addition shows increased affinity for sections with higher protofilament numbers. During copolymerization of the isotypes into a mixed-isotype microtubules, we observe formation of regions with distinct isotype composition and protofilament number, which are then specifically recognized by TPPP1 and OPTN. In the structured part of β-tubulin, most of the surface-exposed amino acid differences between α1β4 and α1β3 are concentrated on the surface that is concealed between the protofilaments. These differences affect the taxol binding pocket, resulting in differences in taxol sensitivity of the isotypes (Chew & Cross, 2023). Surface-exposed amino acid sequence differences between α1β4 and α1β3 accessible in polymerised microtubules are focused in the unstructured C-terminal tails of the molecules. These differences might contribute the differential sensitivity of the MAPs of our interest to the isotypes.

Sectioning of the microtubule lattice during co-polymerization of tubulin isotypes might be a consequence of differences in the dynamic behaviour of the isotypes during microtubule polymerization. In single-isotype microtubules, the most striking differences observed are that α1β4 is less dynamic exhibiting fewer catastrophe and rescue events as compared to α1β3 (Chew & Cross, 2023). Some amino acid differences in the lateral interfaces of β3- and β4-tubulin (specifically the H1-S2 loop and H2-S3 loop) could be the reason for the increased dynamic behaviour of α1β3 microtubules (Wood & Moore, 2025). These differences could potentially prioritise lateral interactions between tubulin dimers of the same isotype composition thereby resulting in the observed tubulin isotype “sorting” in dynamically growing microtubules.

By polymerizing microtubules from HeLa tubulin, a native mixture of different tubulin isotypes (Banerjee et al., 1988) that mostly lacks post-translational modifications, we were able to see the formation of sections with a clearly higher affinity for OPTN and TPPP1, demonstrating that native mixtures of tubulin isotypes assemble into sectioned microtubule lattices, and that this emergent sorting occurs independent of tubulin post-translational modifications. As the predominant β-tubulin isotypes in HeLa microtubules are β4 and β5, while β3 is absent (Banerjee et al., 1988), this suggests that our observation of isotype sorting made with recombinant β3- and β4-tubulin is also relevant for other tubulin isotypes. Microtubules polymerised from brain tubulin also showed sectioning as deduced from patchy OPTN or TPPP1 binding, however the sectioning was less pronounced as for HeLa microtubules. Brain tubulin contains 58% β2, 25% β3, and 13% β4 (Banerjee et al., 1988) and is highly post-translationally modified, which may add further layers of complexity and potentially interfering with the effect of tubulin isotypes.

Protofilament numbers of microtubules assembled *in vitro* from mixed-isotype tubulin can range from 11 to 16, depending on the conditions during polymerization (Andreu et al., 1994). By contrast, *in-vitro* assembled single-isotype microtubules seem to strictly control protofilament numbers regardless of the conditions the microtubules are polymerized and stabilized. We found here that α1β4 microtubules have predominantly 14 protofilaments, while those made of α1β3 have mostly 13. Electron microscopy studies (Chrétien et al., 1992; Guyomar et al., 2022) have reported switches in protofilament numbers during polymerization of mixed-isotype microtubules. This observation aligns with our findings of switching of protofilament numbers during isotype co-polymerization, which we were able to visualize by changes in the signal intensity in IRM, and secondarily by the differential binding of TPPP1 or OPTN. Our results are obtained by *in vitro* studies, thus whether similar processes take place in cells remains to be determined. Given that microtubules with distinct protofilament numbers have been reported in certain cell types (Raff et al., 1997; Renauld et al., 2021; Savage et al., 1989), it is likely that the here-identified isotype sorting does also happen in cells. Changes in protofilament numbers might be mitigated by microtubule interactors affecting these numbers, such as doublecortin (DCX)(Bechstedt & Brouhard, 2012).

In summary, our work sheds light on the organization of the microtubule lattice generated by copolymerizing mixtures of tubulin isotypes. Contrary to the traditional view that polymerization of mixed tubulin isotypes generates a uniformly isotype-mixed lattices (Lewis et al., 1987) we demonstrate that tubulin isotypes can organize the lattice into sections, and that these sections differ in their structural and biological properties. We thus provide the first experimental proof for a hypothesis formulated almost half a century ago (Fulton, 2022): that microtubules assembled from distinct isotypes can become functionally distinct. The evidence of isotype sorting during microtubule polymerisation we provide here now demonstrates that it will be possible to generate distinct microtubule populations in cells that co-exist side-by-side and that fulfil different functions by interacting with distinct effector proteins.

## Methods

### Production of recombinant proteins

N-terminally histidine- and mNeonGreen-tagged full length human optineurin (mNG-OPTN) was expressed and purified according to (Liu et al., 2025). CKAP5 was expressed and purified according to former publication (Sabo et al., 2024).

Human recombinant tubulin isotype α1β4 and α1β3 were produced in Rob Cross lab, of two human β4 isotypes β4a and β4b, we used β4b, herein referred to as β4 (Chew & Cross, 2023), and HeLa tubulin was produced in Carsten Janke lab (Souphron et al., 2019). Tubulin is stored in KPEM buffer (100 mM PIPES, 2 mM EGTA, 1 mM MgSO_4_, pH adjusted to 6.9 with KOH pellets) in the vapour phase of a liquid nitrogen Dewar. The aliquots were retrieved and quickly thawed between hands right before use. The isotypes α1β4 is labelled using HIS lite iFluor 647 Tris NTA chelator (https://www.aatbio.com/products/his-lite-ifluor-647-tris-nta-chelator?unit=12658) for visualization. The dye was reconstituted in MilliQ at a final concentration of 50 μM. 1/10 of tubulin volume of the dye was added to the tubulin so the final concentrations of the α1β4 and dye are about 31.8 μM and 4.5 μM. The reactions were incubated on ice for 15 mins. During the incubation time, the spin columns were buffer exchanged into KPM buffer without EGTA to prevent chelation with the dye. Unbound dyes were removed by the spin columns. The Micro Bio-gel P30 spin columns (Bio-rad, 7326250, come in 10 mM Tris buffer, pH 7.4) were buffer exchanged according to manufacturer’s protocol at 4°C. The final tubulin labelled concentrations were estimated to be about 25 uM.

The pTRIP-CAG-GFP-spacer-hTPPP1 (Jijumon et al., 2022) was expressed in Human embryonic kidney cells (HEK). Cells are seeded on 25 cm2 tissue culture flasks, 2 flasks were used (2 flasks for sufficient quantity of lysate) in growth medium (DMEM + 10% FBS + PENSTREP + 2mM L-glutamine). Cells are transfected using 10 µl of jet PEI reagent per flask, transfection medium per flask: 5 ug of DNA (plasmid), 10 µl of jetPEI (polyplus), DMEM + 10% FBS + PENSTREP + 2mM L-glutamine. The harvesting and cell lysate preparation of over expressed GFP-hTPPP1 was done according to Jijumon et al. 2022.

### *In vitro* reconstitution motility assays

Microtubules were polymerized from HeLa tubulin or recombinant human tubulin or pig brain tubulin in the BRB80 (80 mM PIPES, 1 mM EGTA, 1 mM MgCl_2_, pH 6.9) supplemented with 1 mM MgCl_2_ and 1 mM GTP (Jena Bioscience, Jena, Germany) for 30 minutes at 37 °C prior to centrifugation at 18.000 x g for 30 min in a Microfuge 18 Centrifuge (Beckman Coulter) and resuspension of the microtubule pellet in BRB80 supplemented with 10 µM taxol (Sigma Aldrich, T7191) (BRB80T). GDP microtubules were polymerized the same way except stabilization was done using BRB80 buffer with 40 % Glycerol.

To prepare GMPCPP stabilized or 14 protofilament microtubules, tubulin was incubated with GMPCPP, 1 mM MgCl_2_ in BRB80 for 2 hours at 37 °C prior to centrifugation at 18.000 x g for 30 minutes in a centrifuge (Beckman Coulter) and resuspension of the microtubule pellet in BRB80.

To prepare 12 protofilament microtubules, brain tubulin at a concentration of 40 uM was incubated with sodium phosphate buffer at pH 7, 1 mM MgCl_2_, 1 mM GTP, 2 uM Taxol, and polymerized for 30 minutes at 37 °C. Later the sample was centrifuged at 18.000 x g for 30 minutes. The pellet was resuspended in BRB80 buffer with 10 uM Taxol.

To prepare mixed isotype microtubules, a mixture of α1β3 and unlabelled/iFlour-647 α1β4 was mixed in 1:2 concentration ratio prior to adding polymerization mix (BRB80 supplemented with 1 mM MgCl_2_ and 1 mM GTP) and incubated on ice for 5 minutes. Later polymerization was done at 37 °C for 30 minutes and centrifugation done at 18.000 x g for 30 min in a Microfuge 18 Centrifuge and resuspension of the microtubule pellet in BRB80 supplemented with 10 µM taxol.

Flow cells were prepared by attaching two cleaned and silanized (by DDS (0.05% dichloro-dimethyl silane in trichloroethylene) or HMDS (Bis(trimethylsilyl)amine) glass coverslips (22 x 22 mm^2^ and 18 x 18 mm^2^; Corning, Inc., Corning, NY, USA) together using heated strips of parafilm M (Pechiney Plastic Packaging, Chicago, IL, USA). The flow cells were incubated with 20 µg/ml anti- β-tubulin antibody (Sigma Aldrich,) or a combination of anti- β-tubulin and anti- α- tubulin antibody (produced in Carsten Janke lab) in PBS for 10 min and passivated by 1% pluronic F-127 (Sigma Aldrich, P2443) in PBS for at least 1 hour. The flow cells were then washed with BRB80T, microtubules were introduced into the channel and immobilized on the antibodies, and unbound microtubules were removed by a flush of BRB80T(for taxol stabilized)/BRB80 (for GMPCPP microtubules). Immediately prior to the experiment, the solution was exchanged by motility buffer (BRB10 containing, 10 mM dithiothreitol, 20 mM d-glucose, 0.1% Tween-20, 0.5 mg/ml casein, 1 mM Mg-ATP, 0.22 mg/ml glucose oxidase, and 20 µg/ml catalase) for 76 nM OPTN and BRB80 buffer with 10 uM taxol for 200X dilution of TPPP1 lysate. For experiments on GDP microtubules, instead of taxol, 40% glycerol was used in motility buffer, for experiments with GMPCPP microtubules, neither taxol nor glycerol was used.

To perform experiments with both OPTN and TPPP1 in the same channel, one MAP was introduced first onto the microtubules, and later washed out with buffer (BRB10 for OPTN and BRB80 buffer supplemented with 100 mM salt for TPPP1), and the second MAP was introduced to the channel. All fluorescent imaging was performed using total internal reflection fluorescence (TIRF) microscopy using an inverted NIKON microscope equipped with Apo TIRF 60x Oil, NA 1.49,WD 0.12 mm objective and PRIME BSI (Teledyne Photometrics) camera or CMOS camera (sCMOS ORCA 4.0 V2, Hamamatsu Photonics) or Nikon ECLIPSE Ti2 with Ring TIRF and FRAP adjustment with Nikon Apo TIRF 60x Oil, NA 1.49, WD 0.12 mm and CMOS camera. The imaging setup was controlled by NIS Elements software (Nikon). Microtubules were imaged using Interference Reflection Microscopy (IRM). Movies were taken with an exposure time of 200 ms or 100 ms over the span of 1 minute with switching between IRM mode and TIRF mode to parallelly image microtubules and fluorescent protein.

To perform MAP binding assay with two types of microtubules, one type of microtubules were first added to the channel, unbound microtubules were washed away with BRB80T and an IRM image was acquired and position was saved in NIS software, later second set of microtubules were flushed in without any change to the frame of imaging and an IRM image was acquired. Later the OPTN/TPPP1 was flushed in the channel.

FRAP experiment was carried out H-Ring TIRF at a magnification of 1x or 1.5x using 60x TIRF oil objective. The stimulation of FRAP was carried out using stimulation line at selected ROIs with 50% laser power while imaging done at 50 ms exposure. The sample was imaged at a frame rate of 1 frame per 0.05 seconds. Before bleaching the sample was imaged for 5 seconds, and after bleaching the sample was imaged for 10 seconds.

### Cryo-sample preparation

A sample of 4 µL of polymerized microtubules was applied to a glow discharged Quantifoil cryoEM grid (R 2/1, 300mesh) and incubated for 10 s in the Leica GP2 Plunge Freezer at room temperature and 80% relative humidity. The grid was then automatically blotted using filter paper for 4 s and frozen by immersion into the liquid ethane cooled down to −180 °C.

### Cryo-EM data collection

Cryo-EM data were acquired in a semi-automated mode using SerialEM software (Mastronarde, 2005) on a Jeol JEM-2100Plus transmission electron microscope (TEM) equipped with a LaB_6_ electron gun and TVIPS XF416 CMOS camera, operated at 200kV. The data were collected at a pixel size of 1.44 Å/px, with an exposure time of 4.0 s (33 frames per image), and a total electron dose of 50 e/Å^2^ per micrograph. The nominal defocus range was set from −2.0 to −2.8 µm. In total, each dataset consisted of more than 250 micrographs.

### Cryo EM data processing

The datasets were processed using CryoSPARC v4 (Punjani et al., 2017) to obtain rough estimations of the 3D models. The datasets were imported into CryoSPARC as movies and corrected for motion artifacts acquired during data collection using the Patch Motion Correction job. The contrast transfer function (CTF) parameters were estimated with the Patch CTF job. Micrographs exhibiting good-quality CTF fits and lacking crystalline ice were selected for the Manual Picker job to pick particles along individual microtubules. Approximately 400 manually picked particles were used for initial 2D classification, and selected well-defined classes were imported into the Filament Tracer job. After visual inspection of particle picks using the Inspect Particle Picks job, particles were extracted from the micrographs with an extraction box size of 400 pixels and subjected to a second round of 2D classification. The selected 2D classes were then used in the Helical Refinement job to generate rough 3D models.

For additional statistical analysis of microtubule width, 45 representative microtubules were selected from each dataset and analysed for average width using ImageJ/FIJI. The analysis employed a macro created by the user alpha2zee, which was downloaded from the image.sc forum (https://forum.image.sc/t/imagej-macro-to-measure-distance-between-two-user-drawn-lines-edges-version-2/82226). Briefly, two segmented lines were drawn along the edges of each microtubule, and the average width of each microtubule was calculated from 20 randomly distributed measurements between the lines. The minimum length of each microtubule segment used for the analysis was 300 nm.

### Image Analysis – in vitro reconstitution experiments

All movies were analysed manually using FIJI software. Kymographs were generated using the FIJI Kymograph Builder. To estimate the density of OPTN/TPPP1 on microtubules for purified proteins or in a lysate, the maximum intensity projection was drawn from the movie, a line was drawn on the microtubule with linewidth of 5 and the fluorescent signal and the area on the whole microtubule was measured and for the background. Density is measured as Raw intensity divided by the area. For normalization, for each experimental sets, the median density was measured for one parameter and all the densities for all parameters were divided by it (parameter chosen mentioned in the figure captions). The densities for high MAP binding sections and low MAP binding-sections on mixed isotype microtubules were measured likewise. To perform background subtraction in IRM mode was done similar to (Mahamdeh & Howard, 2019) to study the protofilament numbers. Short time loops (30 frames) were collected from IRM mode from 5 positions in each channel before flushing of microtubules. Once the microtubules attach, short time loops in IRM mode (30 frames) from 5 positions. The median of the time loops without the microtubules were subtracted using Process>Image calculator>Subtract in FIJI (Schindelin et al., 2012). The background subtracted images were used to measure the IRM signal of the microtubules. To measure the relative percentages of MAP density on the high OPTN/TPPP1 and low sections on mixed isotype microtubules, we measure the MAP density on α1β4 as 100% and calculate the percentage increase or decrease of MAP density on high density and low-density sections on the mixed isotype microtubules.

To measure the recovery time for FRAP, cftool in MATLAB was used to fit and exponential curve against time on x-axis and fluorescent density on the y-axis. From the equation -a*exp(-b*x)+c, and recovery time is measured as 1/b. To fit the probability density curve from the width of the microtubule measured using Cryo-EM, the MATLAB code was obtained from ChatGPT AI.

Co-relation plot for OPTN density in comparison with iFluor-647-α1β4 was done using MATLAB with plotting OPTN densities on x axis against iFluor-647 densities on the y-axis. Pearson’s coefficient was calculated using the online tool https://www.statskingdom.com/correlation-calculator.html.

#### Statistical analyses

MATLAB was used to generate graphs and PRISM online software was used for statistical t tests analyses.

#### Data and materials availability

All data are available in the main text or the supplementary materials.

## Author contributions

M.P, M.B and Z.L designed all experiments, R.A.C and C.J offered many experimental ideas and suggestion, M.P performed all of the experiments unless mentioned otherwise, including the analysis and purifications of OPTN. The expression and purification of tubulin isotypes and labelling of isotypes was performed by Y.M.C, the analysis for protofilaments numbers CKAP5 assay, assays with GMPCPP, was performed by D.O, J.M performed Cryo-EM study and analysis, A.K purified HeLa tubulin. M.P, M.B, Z.L, R.A.C and C.J. wrote the manuscript. J.M wrote sections for Cryo-EM.

## Acknowledgments

We are thankful to Yang Hu and Dong Liu (Stanford University), for inspiring us to continue our research with OPTN and for OPTN construct. We are grateful to Katerina Konecna (Institute of Biotechnology CAS) for the expression of OPTN and additional technical support, Kristýna Kroftová (Institute of Biotechnology CAS) for technical support, Jan Sabo (Institute of Biotechnology CAS) for providing CKAP5, his valuable insights and excellent ideas for IRM measurements and reading the manuscript, Alexander Beber (Institute of Biotechnology CAS) for assistance with FRAP analysis. We thank Shivani Ramakrishnan (Institut Curie) for her assistance with HeLa tubulin preparation, and Sudarshan Gadadhar (Institut Curie) for providing anti-tubulin antibodies. The authors acknowledge Imaging Methods Core Facility at BIOCEV, for their support and assistance in this work.

## Supplementary data

**Figure S1:**
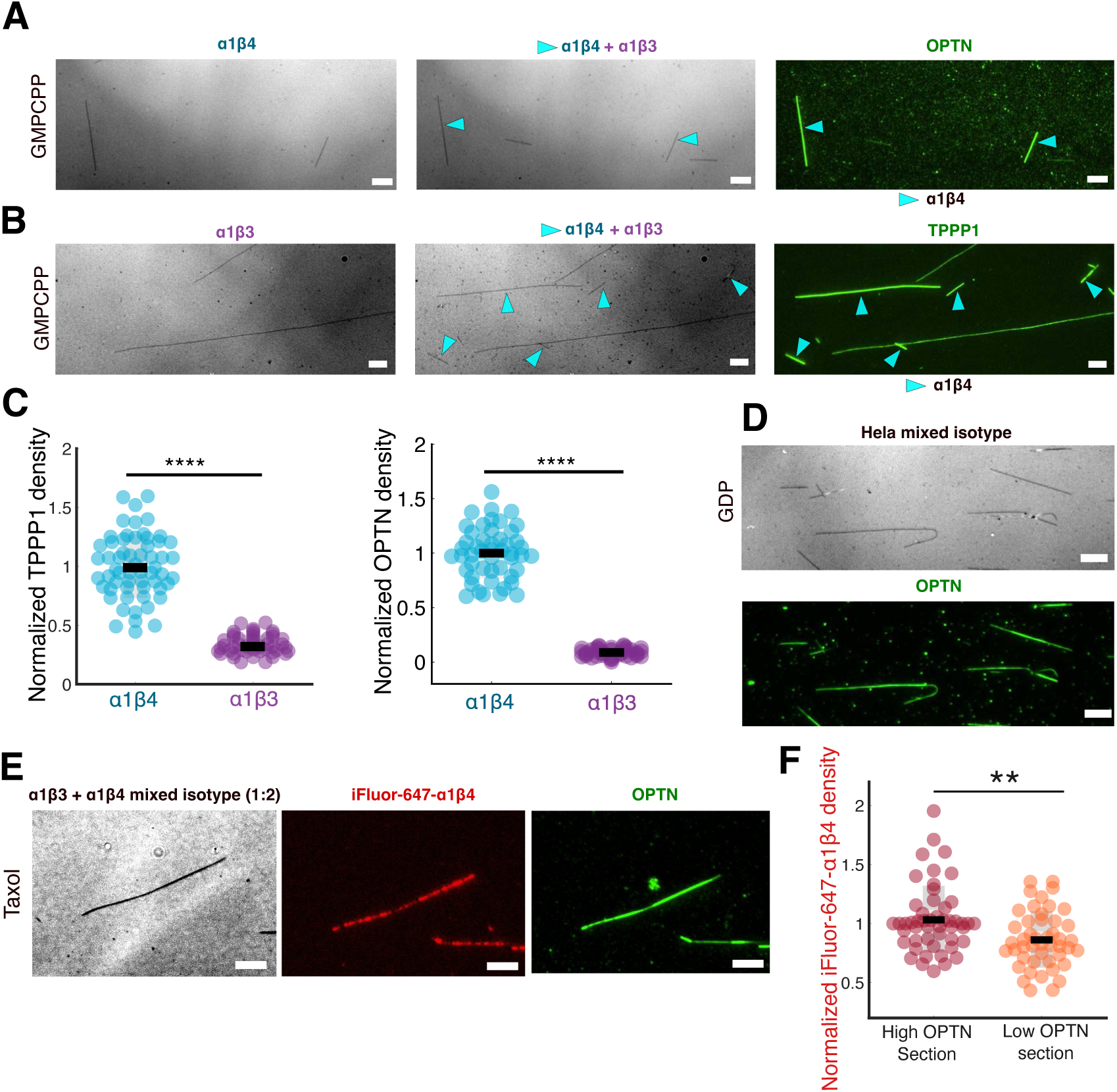
A.B) IRM images of GMPCPP stabilized α1β3 and α1β4 single-isotype-microtubules are sequentially flushed in and later A) OPTN and B) TPPP1 flushed in. C) Analysis of (left) TPPP1 and (right) OPTN density on GMPCPP stabilized α1β4 and α1β3 microtubules (Normalized with average measure), D) (Top to bottom) IRM of HeLa GDP microtubule. OPTN on HeLa GDP microtubules recognizing high/low density sections. E) (left to right) IRM image of mixed isotype microtubules with α1β4 labelled with iFluor-647-tag copolymerized with α1β3 in 1:2 ratio of α1β3 to α1β4, iFluor-647 channel, and OPTN on the microtubule. F) Analysis of iFluor-647 density measured on the high OPTN section and low OPTN section, p value is 0.0022. Scale bar is 4 μm. **** means p<0.0001. **p<0.01. N= 3 for each experiment in figure. Student’s t-test in each plot in figure.

**Methods Figure S2:**
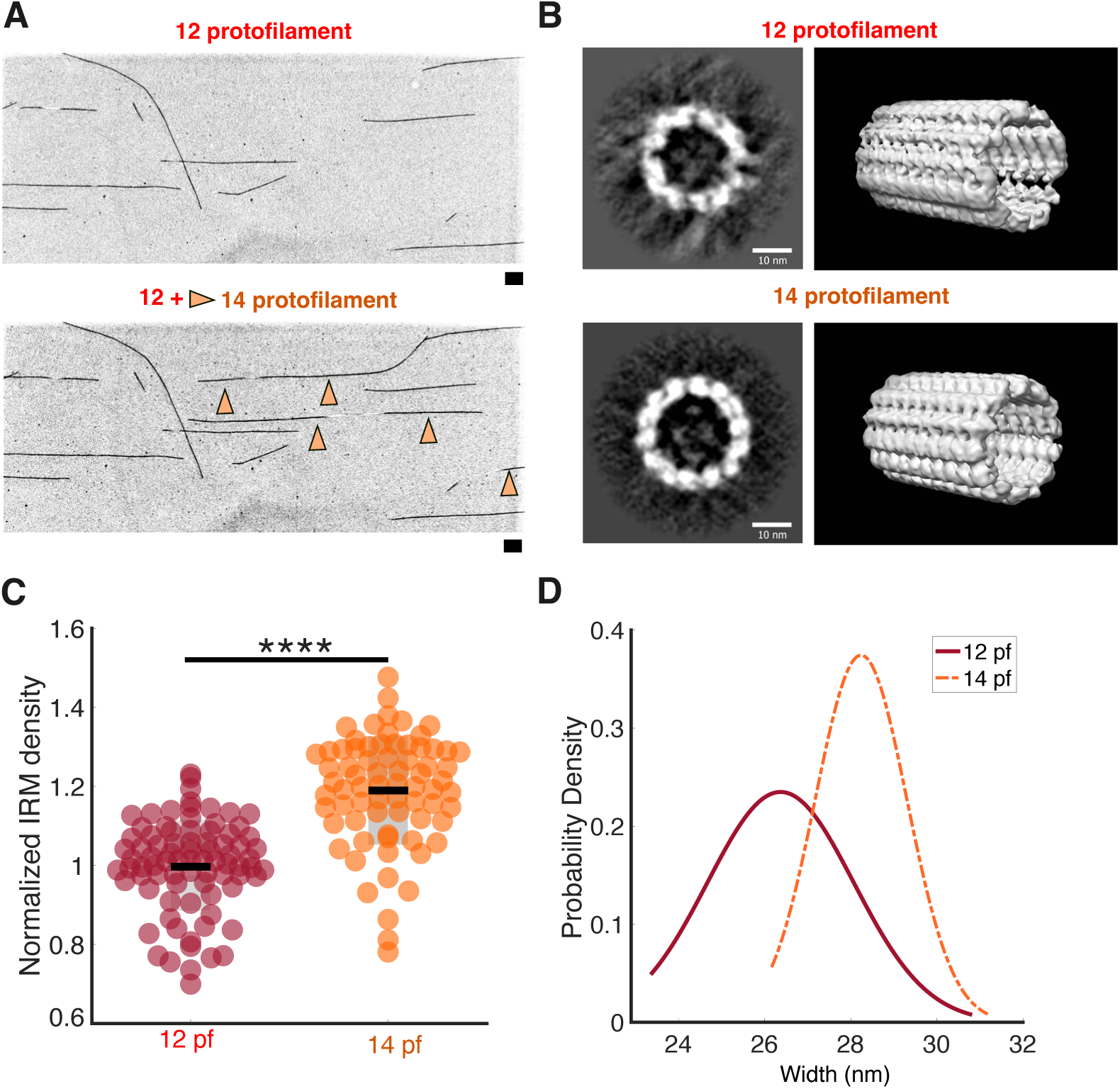
IRM approach to quantifying microtubule protofilament numbers. A) (top to bottom) IRM images of sequential flushes of 12 protofilaments and 14 protofilaments microtubules, orange arrows points to 14 protofilament microtubules. Scale bar is 4 μm. B) Cross-sectional view (left, scale bar 10 μm) and longitudinal view (right) of the 3D model reconstructed from ciyo-EM data. C) IRM quantification displaying the difference in protofilament numbers of 12 and 14 protofilament microtubules (Nonnalized using average), p<0.0001. N=4. D) Probability density of microtubule diameter for 12 and 14 protofilament microtubules measured from Cryo EM images. Student’s t-test in each plot in figure.

**Figure S3:**
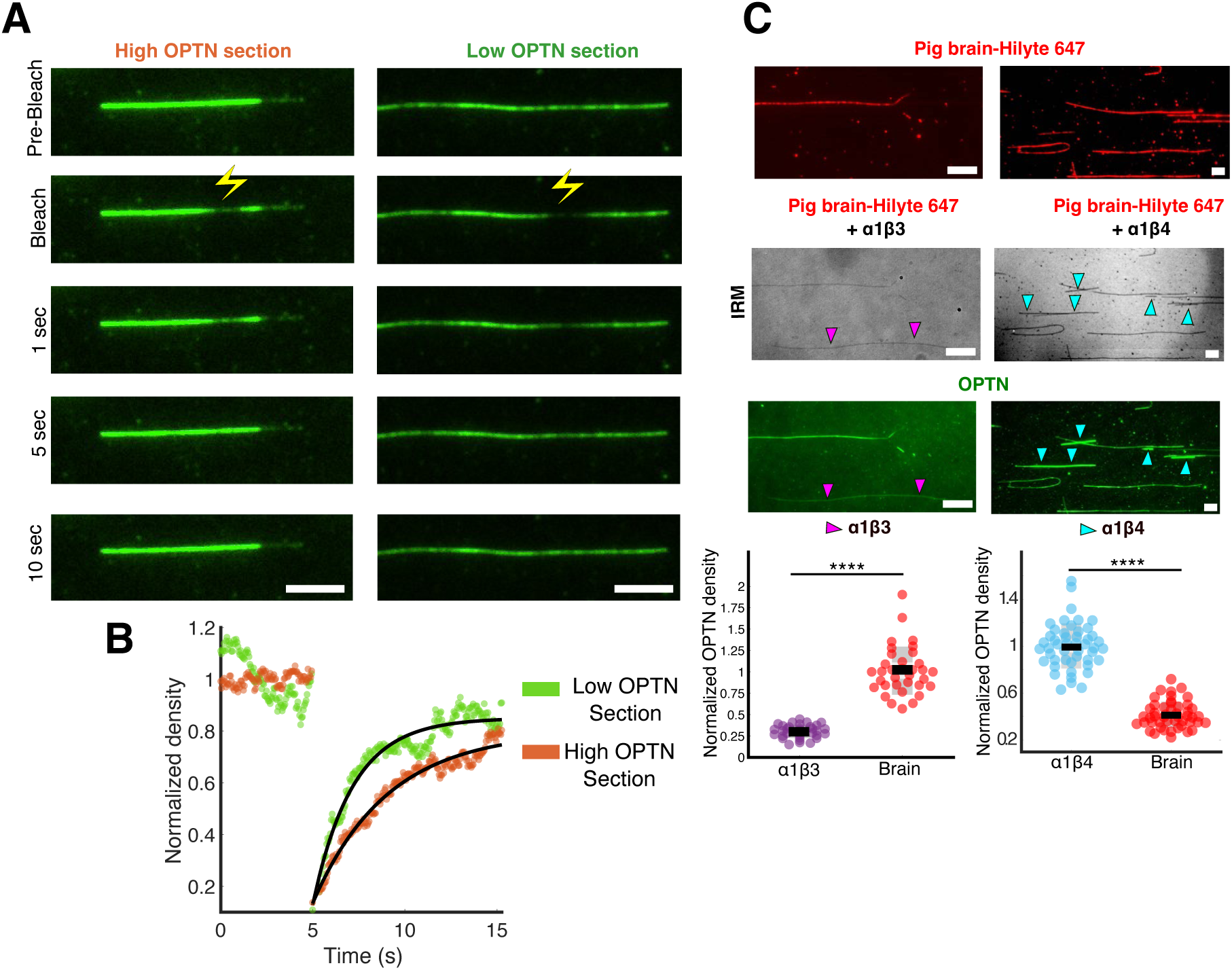
A) (Top to bottom) FRAP on high OPTN sections and low OPTN section yellow lightning bolt indicates the position of bleaching, B) Representative recovery curves of FRAP for high OPTN section (orange) and low OPTN section (green) from (A), recovery time as 2.18 seconds for the low OPTN-section and 4.05 for high-OPTN-section. C) Labelled pig brain microtubules with 647 tag flushed in. position denoted by TIRF microscopy, later label flee polymerized isotypes - α1β4b/- α1β3 is flushed in sequentially and imaged using IRM microscopy. Later. OPTN is flushed in and microtubule binding is recorded and density of OPTN on microtubules is analyzed (Normalization using average). Horizontal scale bar is is 4 μm. **** means p<0.0001, N= 3 for each plot.

## References

Andreu, J. M., Díaz, J. F., Gil, R., de Pereda, J. M., García de Lacoba, M., Peyrot, V., Briand, C., Towns-Andrews, E., & Bordas, J. (1994). Solution structure of Taxotere-induced microtubules to 3-nm resolution. The change in protofilament number is linked to the binding of the taxol side chain. The Journal of Biological Chemistry, 269(50), 31785– 31792.

Banerjee, A., Roach, M. C., Wall, K. A., Lopata, M. A., Cleveland, D. W., & Ludueña, R. F. (1988). A monoclonal antibody against the type II isotype of beta-tubulin. Preparation of isotypically altered tubulin. The Journal of Biological Chemistry, 263(6), 3029–3034.

Barisic, M., Silva e Sousa, R., Tripathy, S. K., Magiera, M. M., Zaytsev, A. V., Pereira, A. L., Janke, C., Grishchuk, E. L., & Maiato, H. (2015). Mitosis. Microtubule detyrosination guides chromosomes during mitosis. Science (New York, N.Y.), 348(6236), 799–803. 10.1126/science.aaa5175

Bechstedt, S., & Brouhard, G. J. (2012). Doublecortin recognizes the 13-protofilament microtubule cooperatively and tracks microtubule ends. Developmental Cell, 23(1), 181–192. 10.1016/j.devcel.2012.05.006

Chaaban, S., & Brouhard, G. J. (2017). A microtubule bestiary: Structural diversity in tubulin polymers. Molecular Biology of the Cell, 28(22), 2924–2931. 10.1091/mbc.E16-05-0271

Chew, Y. M., & Cross, R. A. (2023). Taxol acts differently on different tubulin isotypes. Communications Biology, 6(1), 946. 10.1038/s42003-023-05306-y

Chrétien, D., Metoz, F., Verde, F., Karsenti, E., & Wade, R. H. (1992). Lattice defects in microtubules: Protofilament numbers vary within individual microtubules. The Journal of Cell Biology, 117(5), 1031–1040. 10.1083/jcb.117.5.1031

Fulton, C. (2022). The Amazing Evolutionary Complexity of Eukaryotic Tubulins: Lessons from Naegleria and the Multi-tubulin Hypothesis. Frontiers in Cell and Developmental Biology, 10, 867374. 10.3389/fcell.2022.867374

Guyomar, C., Bousquet, C., Ku, S., Heumann, J. M., Guilloux, G., Gaillard, N., Heichette, C., Duchesne, L., Steinmetz, M. O., Gibeaux, R., & Chrétien, D. (2022). Changes in seam number and location induce holes within microtubules assembled from porcine brain tubulin and in Xenopus egg cytoplasmic extracts. eLife, 11, e83021. 10.7554/eLife.83021

Hyman, A. A., Chrétien, D., Arnal, I., & Wade, R. H. (1995). Structural changes accompanying GTP hydrolysis in microtubules: Information from a slowly hydrolyzable analogue guanylyl-(alpha,beta)-methylene-diphosphonate. The Journal of Cell Biology, 128(1–2), 117–125. 10.1083/jcb.128.1.117

Janke, C., & Magiera, M. M. (2020). The tubulin code and its role in controlling microtubule properties and functions. Nature Reviews. Molecular Cell Biology, 21(6), 307–326. 10.1038/s41580-020-0214-3

Jijumon, A. S., Bodakuntla, S., Genova, M., Bangera, M., Sackett, V., Besse, L., Maksut, F., Henriot, V., Magiera, M. M., Sirajuddin, M., & Janke, C. (2022). Lysate-based pipeline to characterize microtubule-associated proteins uncovers unique microtubule behaviours. Nature Cell Biology, 24(2), 253–267. 10.1038/s41556-021-00825-4

Kalutskii, M., Grubmüller, H., Volkov, V. A., & Igaev, M. (2024). Microtubule dynamics are defined by conformations and stability of clustered protofilaments. bioRxiv, 2024.11.04.621893. 10.1101/2024.11.04.621893

Keates, R. A. (1980). Effects of glycerol on microtubule polymerization kinetics. Biochemical and Biophysical Research Communications, 97(3), 1163–1169. 10.1016/0006-291x(80)91497-7

Kikuchi, T., Takasaka, T., Tonosaki, A., Katori, Y., & Shinkawa, H. (1991). Microtubules of guinea pig cochlear epithelial cells. Acta Oto-Laryngologica, 111(2), 286–290. 10.3109/00016489109137389

Kikuchi, T., Takasaka, T., Tonosaki, A., Watanabe, H., Hozawa, K., Shinkawa, H., & Wada, H. (1991). Microtubule subunits of guinea pig vestibular epithelial cells. Acta Oto-Laryngologica. Supplementum, 481, 107–111. 10.3109/00016489109131359

Lewis, S. A., Gu, W., & Cowan, N. J. (1987). Free intermingling of mammalian beta-tubulin isotypes among functionally distinct microtubules. Cell, 49(4), 539–548. 10.1016/0092-8674(87)90456-9

Liu, D., Webber, H. C., Bian, F., Xu, Y., Prakash, M., Feng, X., Yang, M., Yang, H., You, I.-J., Li, L., Liu, L., Liu, P., Huang, H., Chang, C.-Y., Liu, L., Shah, S. H., La Torre, A., Welsbie, D. S., Sun, Y., … Hu, Y. (2025). Optineurin-facilitated axonal mitochondria delivery promotes neuroprotection and axon regeneration. Nature Communications, 16(1), 1789. 10.1038/s41467-025-57135-8

Mahamdeh, M., & Howard, J. (2019). Implementation of Interference Reflection Microscopy for Label-free, High-speed Imaging of Microtubules. Journal of Visualized Experiments: JoVE, 150. 10.3791/59520

Mastronarde, D. N. (2005). Automated electron microscope tomography using robust prediction of specimen movements. Journal of Structural Biology, 152(1), 36–51. 10.1016/j.jsb.2005.07.007

Mitchison, T., & Kirschner, M. (1984). Dynamic instability of microtubule growth. Nature, 312(5991), 237–242. 10.1038/312237a0

Nogales, E. (2000). Structural insights into microtubule function. Annual Review of Biochemistry, 69, 277–302. 10.1146/annurev.biochem.69.1.277

Nsamba, E. T., & Gupta, M. L. (2022). Tubulin isotypes—Functional insights from model organisms. Journal of Cell Science, 135(9), jcs259539. 10.1242/jcs.259539

Punjani, A., Rubinstein, J. L., Fleet, D. J., & Brubaker, M. A. (2017). cryoSPARC: Algorithms for rapid unsupervised cryo-EM structure determination. Nature Methods, 14(3), 290–296. 10.1038/nmeth.4169

Raff, E. C., Fackenthal, J. D., Hutchens, J. A., Hoyle, H. D., & Turner, F. R. (1997). Microtubule architecture specified by a beta-tubulin isoform. *Science (New York*, N.Y.), 275(5296), 70–73. 10.1126/science.275.5296.70

Ray, S., Meyhöfer, E., Milligan, R. A., & Howard, J. (1993). Kinesin follows the microtubule’s protofilament axis. The Journal of Cell Biology, 121(5), 1083–1093. 10.1083/jcb.121.5.1083

Renauld, J., Thelen, N., Bartholomé, O., Malgrange, B., & Thiry, M. (2021). Dispensability of Tubulin Acetylation for 15-protofilament Microtubule Formation in the Mammalian Cochlea. Cell Structure and Function, 46(1), 11–20. 10.1247/csf.20057

Sabo, J., Dujava Zdimalova, M., Slater, P. G., Dostal, V., Herynek, S., Libusova, L., Lowery, L. A., Braun, M., & Lansky, Z. (2024). CKAP5 enables formation of persistent actin bundles templated by dynamically instable microtubules. Current Biology: CB, 34(2), 260–272.e7. 10.1016/j.cub.2023.11.031

Savage, C., Hamelin, M., Culotti, J. G., Coulson, A., Albertson, D. G., & Chalfie, M. (1989). Mec-7 is a beta-tubulin gene required for the production of 15-protofilament microtubules in Caenorhabditis elegans. Genes & Development, 3(6), 870–881. 10.1101/gad.3.6.870

Schindelin, J., Arganda-Carreras, I., Frise, E., Kaynig, V., Longair, M., Pietzsch, T., Preibisch, S., Rueden, C., Saalfeld, S., Schmid, B., Tinevez, J.-Y., White, D. J., Hartenstein, V., Eliceiri, K., Tomancak, P., & Cardona, A. (2012). Fiji: An open-source platform for biological-image analysis. Nature Methods, 9(7), 676–682. 10.1038/nmeth.2019

Schofield, A., & Bernard, O. (2013). Tubulin polymerization promoting protein 1 (TPPP1). Communicative & Integrative Biology, 6(6), e26316. 10.4161/cib.26316

Souphron, J., Bodakuntla, S., Jijumon, A. S., Lakisic, G., Gautreau, A. M., Janke, C., & Magiera, M. M. (2019). Purification of tubulin with controlled post-translational modifications by polymerization-depolymerization cycles. Nature Protocols, 14(5), 1634–1660. 10.1038/s41596-019-0153-7

Ti, S.-C., Alushin, G. M., & Kapoor, T. M. (2018). Human β-Tubulin Isotypes Can Regulate Microtubule Protofilament Number and Stability. Developmental Cell, 47(2), 175–190.e5. 10.1016/j.devcel.2018.08.014

Tilney, L. G., Bryan, J., Bush, D. J., Fujiwara, K., Mooseker, M. S., Murphy, D. B., & Snyder, D. H. (1973). Microtubules: Evidence for 13 protofilaments. The Journal of Cell Biology, 59(2 Pt 1), 267–275. 10.1083/jcb.59.2.267

Vemu, A., Atherton, J., Spector, J. O., Moores, C. A., & Roll-Mecak, A. (2017). Tubulin isoform composition tunes microtubule dynamics. Molecular Biology of the Cell, 28(25), 3564– 3572. 10.1091/mbc.E17-02-0124

Wood, L. M., & Moore, J. K. (2025). Β3 accelerates microtubule plus end maturation through a divergent lateral interface. Molecular Biology of the Cell, 36(4), ar36. 10.1091/mbc.E24-08-0354

